# How asbestos drives the tissue towards tumors: YAP activation, macrophage and mesothelial precursor recruitment, RNA editing and somatic mutations

**DOI:** 10.1101/182980

**Authors:** Hubert Rehrauer, Licun Wu, Walter Blum, Lazslo Pecze, Thomas Henzi, Véronique Serre-Beinier, Catherine Aquino, Bart Vrugt, Marc de Perrot, Beat Schwaller, Emanuela Felley-Bosco

**Affiliations:** Functional Genomics Center Zurich, ETH Zurich and University of Zurich, 8057 Zurich, Switzerland; Latner Thoracic Surgery Research Laboratories, Division of Thoracic Surgery, Toronto General Hospital, University Health Network, University of Toronto, Toronto, ON Canada; Unit of Anatomy, Department of Medicine, University of Fribourg, Route Albert-Gockel 1, CH-1700 Fribourg, Switzerland; Department of Thoracic Surgery, University Hospitals of Geneva, Geneva, Switzerland; Institute of Surgical Pathology, University Hospital Zurich; Laboratory of Molecular Oncology, Lungen- und Thoraxonkologie Zentrum, University Hospital Zürich, Sternwartstrasse 14, 8091 Zurich, Switzerland

**Keywords:** asbestos, pre-cancer alterations, mesothelium tissue homeostasis, YAP/TAZ activation, chronic inflammation, RNA editing

## Abstract

Chronic exposure to intraperitoneal asbestos triggered a marked response in the mesothelium well before tumor development. Macrophages, mesothelial precursor cells, cytokines and growth factors accumulated in the peritoneal lavage. Transcriptome profiling revealed YAP/TAZ activation in inflamed mesothelium with further activation in tumors, paralleled by increased levels of cells with nuclear YAP/TAZ. *Arg1* was one of the highest upregulated genes in inflamed tissue and tumor. Inflamed tissue showed increased levels of single nucleotide variations, with an RNA-editing signature, which were even higher in the tumor samples. Subcutaneous injection of asbestos-treated, but tumor-free mice with syngeneic mesothelioma tumor cells resulted in a significantly higher incidence of tumor growth when compared to naïve mice supporting the role of the environment in tumor progression.

**Highlights:**

- Asbestos increases levels of cytokines and growth factors in mesothelium environment
- Recruitment of macrophages and mesothelial precursor cells prior to tumor development
- YAP/TAZ signaling upregulated in pre-neoplastic tissues and cancer
- Increased RNA-editing and somatic mutations as early steps in tumor development

## Introduction

Mesothelioma is associated with asbestos exposure and asbestos-related diseases are still a major public health problem (Stayner et al., 2013). The association of exposure to asbestos with development of mesothelioma has been demonstrated in the seminal experimental work of J.C. Wagner in the sixties (Wagner, 1962). In 1987 A. B. Kane and co-workers (Moalli et al., 1987) observed that already a single dose of asbestos fibers damages the mesothelium tissue and stimulates regeneration with the additional recruitment of macrophages to the site of damage. They investigated the “short”-term (few hours until 3 weeks) reaction to acute injury, and were the first to propose that persistent tissue injury leads to an inflammatory and regenerative response, which subsequently paves the way to mesothelioma development. Kane *et al.* also observed that repeated exposure to asbestos leads to angiogenesis surrounding the regenerating mesothelium after six weeks, well before mesothelioma development and suggested that the release of angiogenic factors represents an early biomarker for asbestos exposure (Branchaud et al., 1989).

Tumor development greatly depends on the evasion from the immune surveillance. Tumor cells escaping the immune defense is enhanced by the induction of an immunosuppressive tumor microenvironment (Mapara and Sykes, 2004). Effective adaptive immune responses are suppressed through the activation of several pathways. For example, the differentiation and activation of antigen-presenting dendritic cells, which are the key initiators of the adaptive immune responses (Banchereau and Palucka, 2005), are inhibited by signals such e.g. vascular endothelial growth factor (VEGF), present in the tumor microenvironment (Tartour et al., 2011). Tumor-associated macrophages polarized to a “M2” state are induced in tumor-bearing hosts (Jinushi et al., 2011); these cells, as well as a heterogeneous population of myeloid-derived suppressor cells (MDSCs), are potent suppressors of antitumor immunity. MDSCs and “M2” state macrophages also sustain the malignant behavior of tumor cells by secreting cytokines, growth factors, and proteases, which promote tumor progression or enhance metastasis (Gabrilovich and Nagaraj, 2009). Others and we have previously observed that blockade of immunosuppressive signals enhances immunity against mesothelioma (Cherkassky et al., 2016, Khanna et al., 2016, Veltman et al., 2010, Wu et al., 2011, Wu et al., 2012).

This study assesses the loss of homeostasis in the mesothelial environment during tumor development after exposure to asbestos. Since mesothelioma development after intraperitoneal exposure to asbestos has been widely accepted as a *bona fide* surrogate to investigate mesothelium-dependent reaction, we used this model to investigate in-depth how perturbation of homeostatic control leads to tumor development. Nf2+/- mice on a C57Bl/6J genetic background were used for several reasons including the observation that tumor that develop in Nf2+/- mice exposed to asbestos show common genomic alterations with human mesothelioma and loss of NF2 function has a driver role in mesothelioma development (Jean et al., 2011, Altomare et al., 2005, Fleury-Feith et al., 2003, Jongsma et al., 2008), and moreover C57Bl/6J mice being widely used in functional studies.

## Results

### Exposure to asbestos alters the profile of cell populations and signaling molecules in the peritoneal cavity in non-tumor bearing mice

C57Bl/6J Nf2+/- mice were exposed eight times to crocidolite (blue asbestos) every three weeks. Mice were sacrificed 33 weeks after first crocidolite exposure in order to have the possibility to investigate pre-cancer and cancer stages (Figure 1A). In mesothelioma-free mice, the population of CD68/F480 macrophages in the peritoneal lavage was five-fold higher compared to sham-exposed mice (Figure 1B). This finding is consistent with results of by Moalli *et al.* (Moalli et al., 1987) who showed increased macrophage prevalence a few days after a single-dose of asbestos administration. Both, the numbers of T and B cells (Figure 1B) in the peritoneal lavage of crocidolite-treated mice were significantly decreased.

**Figure 1.**
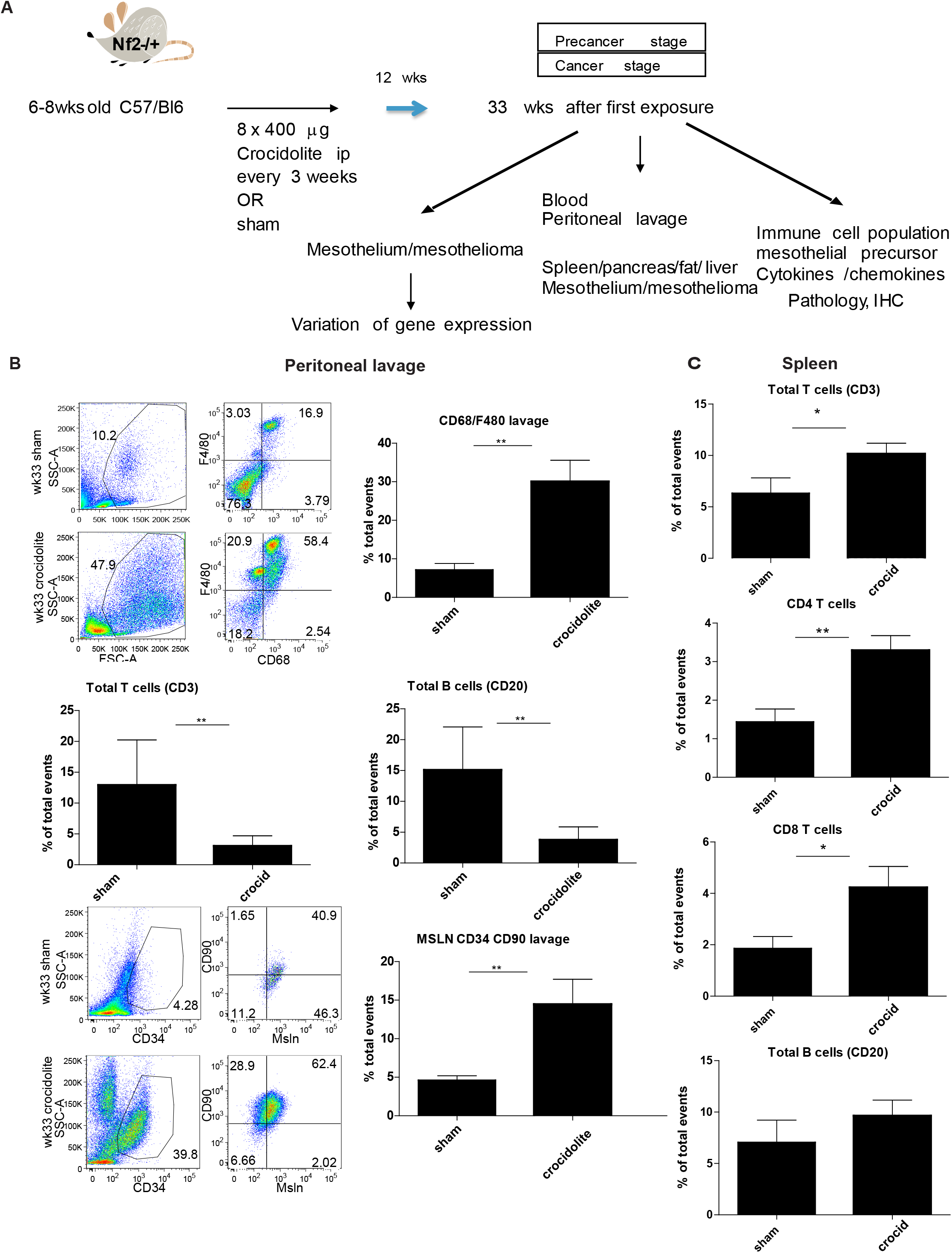
Exposure to asbestos alters the profile of cell populations in peritoneal lavage. **A.** Experimental scheme: 6-8 week-old C57Bl/6J mice were exposed to crocidolite i.p. (400 μg/mouse) every 3 weeks with in total 8 treatments. Thirty-three weeks after initial exposure to crocidolite mice were sacrificed to collect peritoneal lavage, blood and tissues. Tumor bearing mice were sacrificed to collect tumor tissue. **B.** The lavage from crocidolite-exposed mice shows higher proportions of macrophages and mesothelial precursor populations as compared to sham-exposed mice. Flow-cytometry analysis of samples is shown in left panel. A significant decrease in the total number of T and B cells was observed in the peritoneal lavage 12 weeks after last exposure to crocidolite **C.** The proportion of T cells was significantly increased in the spleen of asbestos-exposed mice. No difference for B cells was observed in the spleen. N=6-8 mice. Mean±SE. * p< 0.05, ** p< 0.01, Mann-Whitney test.

Besides macrophage recruitment after tissue damage, free-floating mesothelial-like cells were shown to be incorporated into peritoneal wound surfaces and to contribute to the regeneration of the damaged mesothelium (Foley-Comer et al., 2002). This population is characterized as mesothelin^+^ (Msln^+^)-bone marrow-derived progenitor cells (Carmona et al., 2011). During embryonic development a mesothelial precursor population has been identified as Msln^+^ CD105^low^, CD90^high^, CD44^low^, CD34^high^ (Rinkevich et al., 2012). Accordingly, we observed that CD90^+^CD34^+^Msln^+^ cells paralleled the CD68/F480 cell profile (Figure 1B). In the spleen a significant increase of T cells, both CD4 and CD8, was observed (Figure 1C).

In order to explore mechanisms putatively involved in the observed changes, the content of different inflammatory cytokines (IL-6, IL-10, IFNγ), myeloid chemoattractant chemokines (CCL-2, CCL5, CXCL1) and growth factors modulating the differentiation of myeloid cells (G-CSF, GM-CSF) or vessel growth (VEGF), was measured in the peritoneal lavage (Figure 2). Levels of the above mentioned proteins were increased in crocidolite-exposed mice paralleling the profile change observed for the CD68/F480 and mesothelial precursor cell populations. A strong correlation was observed between CCL2 and IL-6 levels (r=0.7, p<0.0001) consistent with the known reciprocal regulation of these two cytokines (Roca et al., 2009).

**Figure 2.**
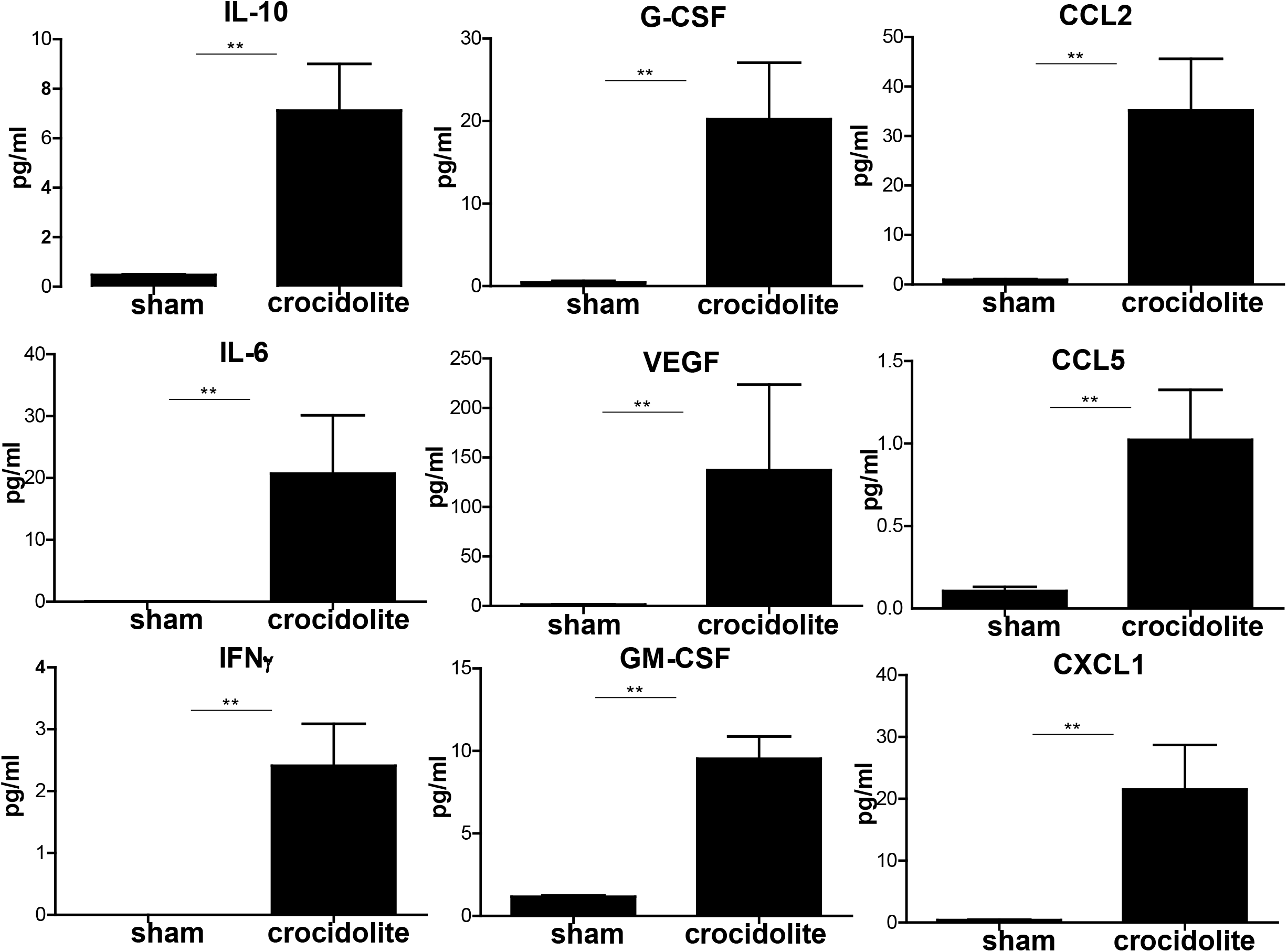
Exposure to asbestos alters the profile of signaling molecules in the peritoneal lavage. 6-8 week-old C57Bl/6J mice were exposed to crocidolite i.p. (400 μg/mouse) every 3 weeks for eight rounds. Thirty-three weeks after initial exposure to crocidolite mice were sacrificed to collect peritoneal lavage to determine the levels of several cytokines, growth factors and chemokines. All of them were more abundant in the lavage from crocidolite-exposed mice. N=6-8 mice. Mean±SE. ** p< 0.01, Mann-Whitney test.

### Mesothelial tissue histopathological changes during mesothelioma development in mice exposed to asbestos

Critical criteria to define mesothelioma were the morphology with high cellularity, cytonuclear atypia, haphazard arrangement of tumor cells and retained cytokeratin expression also in the deep parts of the tumor. Tumors cells growing in crocidolite-exposed mice showed a spindled morphology and stained positive for podoplanin, WT-1 (Figure 3A), supporting mesothelial origin, as well as mesothelin, vimentin and cytokeratin (data not shown). Benign mesothelial proliferations showing similar immunoreactivity were observed on the diaphragm (Figure 3B), serosal surfaces (Figure 3C) and in spheroids implanted on both the visceral (Figure 3D) and parietal (Figure 3E) mesothelial linings.

**Figure 3.**
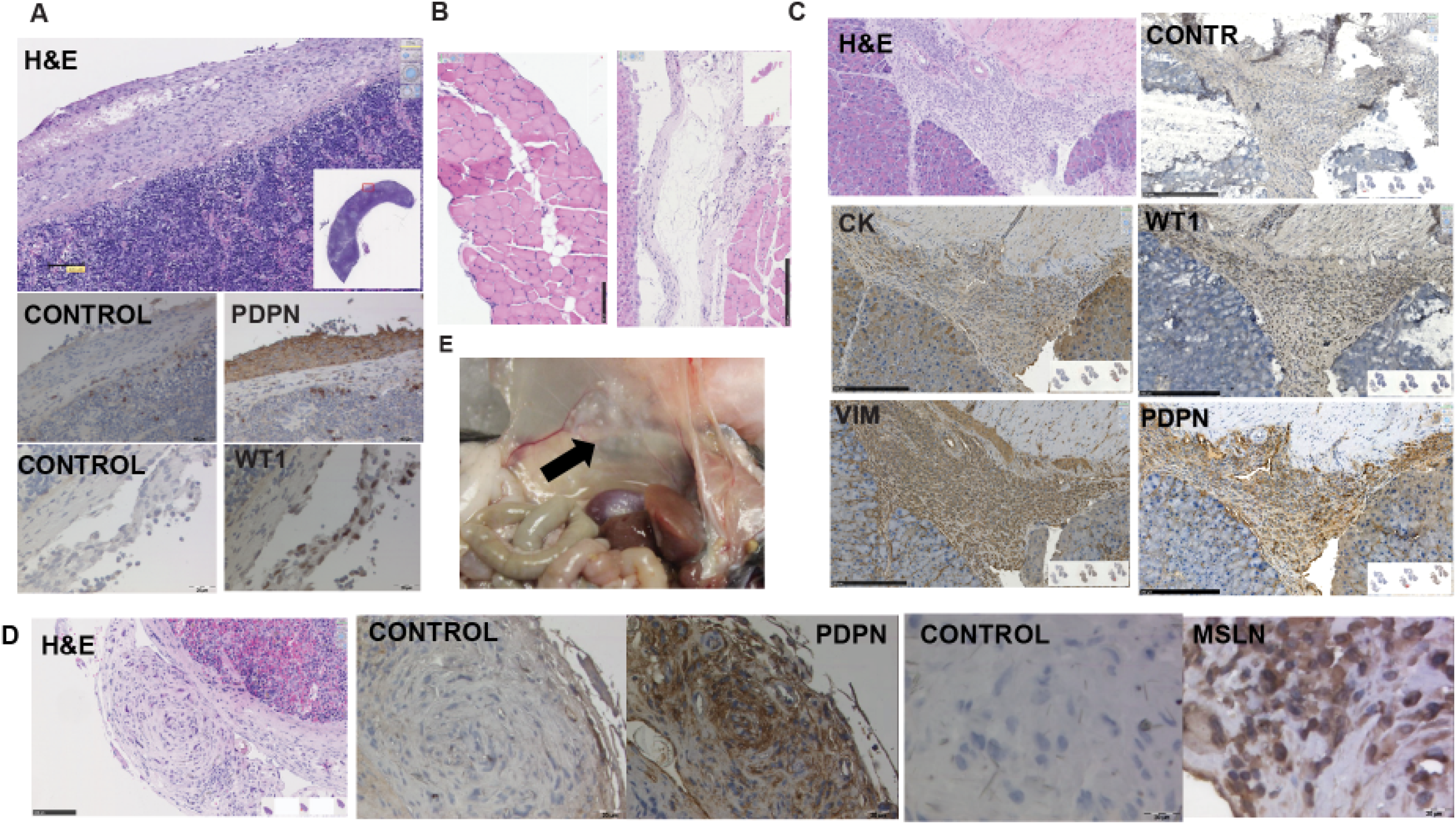
Histopathology of Nf2 (+/-) mice following repeated i.p. injections of crocidolite fibers. **A.** A localized malignant mesothelioma on the surface of the spleen was stained with H&E, podoplanin (PDPN) and WT-1. **B.** Morphology of the diaphragm shows inflammatory cell infiltrates and reactive mesothelium after crocidolite exposure (right panel) compared to the normal single-layer mesothelium in sham-exposed mice (left panel). **C.** Benign mesothelial cell growth on the serosal surface of organs showing co-expression of for vimentin (vim), cytokeratin (CK), WT-1 and PDPN **D.** Benign growth on serosal surfaces and in spheroids implanted on the visceral organs showing immunoreactivity for PDPN and mesothelin (Msln). **E.** Nodules growing after crocidolite exposure on the parietal mesothelium, which was scraped for obtaining tissue for gene expression analysis.

### RNA-seq transcriptome profiling

We collected scraped mesothelium (Figure 4A) and performed RNA-seq to identify gene expression changes during mesotheliomagenesis. We analyzed three treatment groups by RNA-seq: sham, age-matched crocidolite-exposed, and age-matched crocidolite-exposed with observable tumors. Among those groups we performed differential expression analysis between crocidolite-exposed and sham, and detected 5976 differentially expressed genes (p <0.01, FDR < 0.022). Additionally we assessed the expression differences between crocidolite-exposed with tumors and crocidolite-exposed, and identified 8416 genes (p <0.01, FDR <0.017). In Figure 4B we show a heatmap of the significant genes where we applied an additional fold-change threshold of higher than two-fold (2316 genes up, 84 genes down when comparing crocidolite-exposed and sham). The crocidolite response is dominated by an upregulation of genes. It is interesting to note that among the 2316 genes with a positive response, 1989 have an even higher average expression in the crocidolite-with-tumor samples as compared to the crocidolite samples and only 327 genes have a lower average expression. However, this asymmetry is not generally true, when looking at all genes significantly changed between crocidolite with tumors and crocidolite without tumors. We find 3014 genes with more than two-fold upregulation and 3008 genes with a more than two-fold down regulation. 1234 genes of these two upregulated gene sets were overlapping, as shown in the Venn diagram (Figure 4C). The commonly upregulated 1234 genes represent 53% of the upregulated genes in the crocidolite-exposed tissue vs. sham-exposed pool and 41% in the crocidolite-exposed tumor vs crocidolite-inflamed tissue pool (Figure 4C). A functional analysis of these different responding gene groups is discussed below.

**Figure 4.**
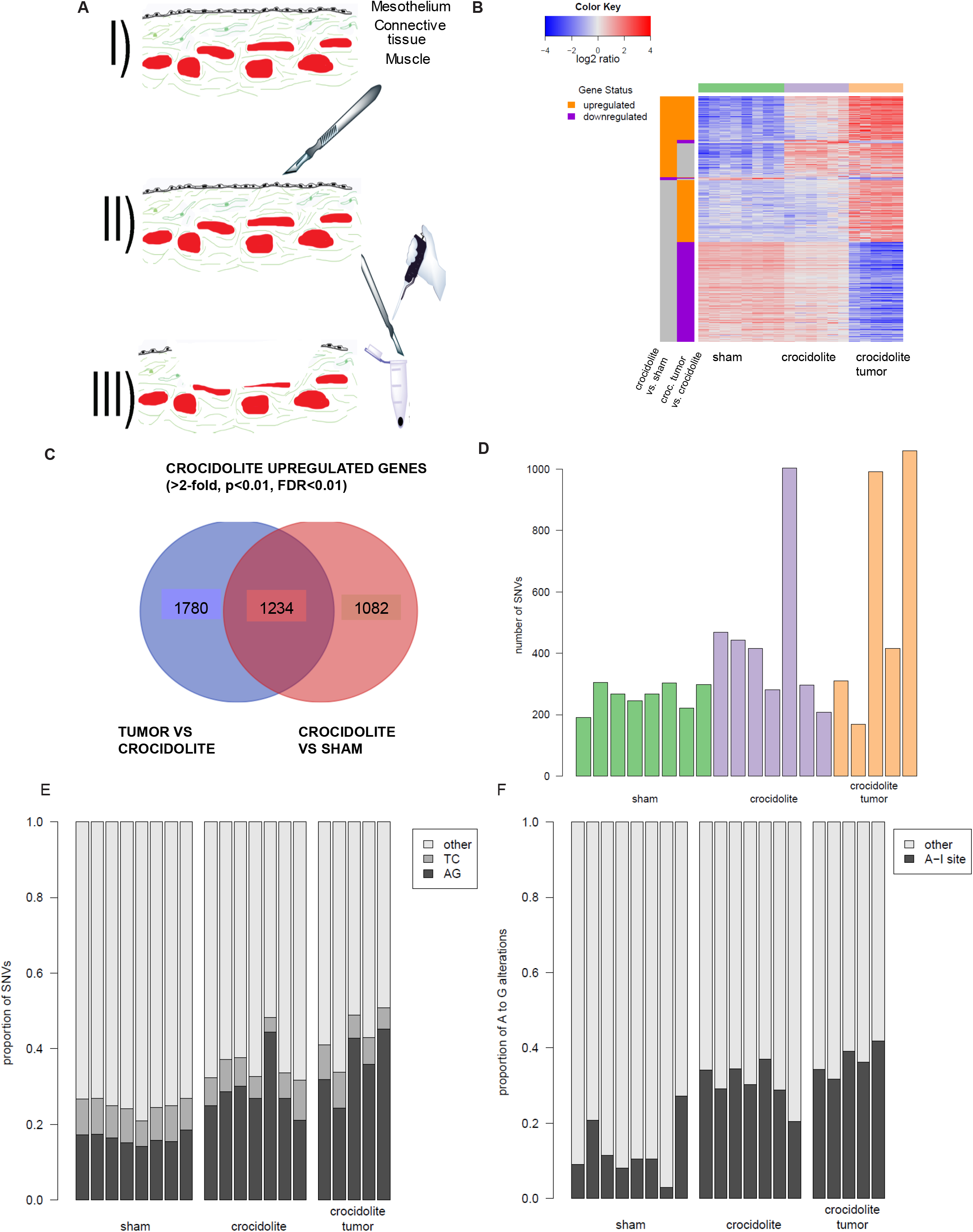
Transcriptome profiles reveal higher expression for thousands of genes and an increase of RNA editing events. **A.** Mesothelial tissue collection by exposure of parietal mesothelium and scraping the surface with a scalpel. **B.** Heatmap of differentially expressed genes in the two comparisons crocidolite vs sham and crocidolite-tumors vs crocidolite. The color bars at the left indicate up-regulation (orange) and down-regulation (violet). The crocidolite vs sham comparison is dominated by genes up-regulated in the crocidolite condition. **C.** Overlap of the differentially expressed genes (2-fold change, p<0.01) in both comparisons visualized as a Venn diagram. There is a substantial overlap that originates from the fact that about 40% of the genes that are upregulated in the crocidolite condition, show an even higher expression in the crocidolite tumor tissue. **D.** Number of detected single nucleotide variations (SNV) in RNA-seq reads. **E.** An increase of A to G transitions is the major cause of the increased number of SNVs in crocidolite-exposed tissues. Please note the reads are strand-specific and the reverse complement transition (T to C) is not increased. **F.** The proportion of A to G mutations that coincide with known A to I RNA-editing sites increases also in the crocidolite exposed tissues.

We further looked at the number and type of base changes in the RNA-seq data. In Figure 4D we show the mutational load as determined from the strand-specific RNA-seq read alignments. The crocidolite-exposed mice show higher number of mutations as compared to sham-treated. After tumorigenesis the number of mutations was even further increased. Figure 4D shows that there is one crocidolite non-tumor sample with a very high number of mutations. This outlier sample has been obtained from a mouse with no visible tumor. In gene expression analysis its profile was intermediate between tumor and crocidolite without tumor samples and histological examination by the pathologist has revealed the presence of some neoplastic cells, which means that it lies in between our two groups “crocidolite non-tumor” and “crocidolite tumor”. Because only one sample exhibited this characteristic, we have excluded it from further expression analysis but included it in the base-change analysis. Excluding it from the base-change analysis does not substantially alter the results. When looking closer at the type of mutations, we observed a significant increase of A to G mutations between crocidolite and sham treated samples. This increase is significant in terms of absolute counts (p=0.005) and in terms of the relative fractions of the A to G mutations of the total mutations (p=0.00031) (Figure 4E). The A to G alterations were increased while the T to C alterations were not, which identifies them as RNA-editing events. Actually, none of the other mutation types showed a significant increase neither in absolute nor in relative terms. Even more, Figure 4F shows that the fraction of A to G alterations that overlaps with known A to I sites (http://rnaedit.com/download/) also increases (p= 0.0012). We hypothesize that they are the result of hydrolytic deamination of adenosine downstream of Adar activity (Nishikura, 2016) (I is detected as G in RNA-seq). This is consistent with a significant 3.9-fold increase of *Adar* expression in inflamed tissue compared to sham (p=5.35E-34, FDR=7.87E-32) and more than two-fold increase in tumors compared to inflamed tissues (p=2.56E-12, FDR=1.86E-11). Intriguingly *Adarb1* showed a significant more than two-fold increase in tumors compared to inflamed tissues (p=2.56E-12, FDR=1.86E-11), but its expression was not significantly changed between sham and inflamed tissue. Analysis of TCGA mesothelioma data revealed that high expression of *ADARB1* is associated with worst overall survival (Figure S1) supporting the idea that RNA editing is relevant in mesothelioma as it is in other cancers.

### Functional analysis of differentially expressed genes

Genes known to be upregulated during mesothelioma development such as *Osteopontin (Spp1)*, and *mesothelin (Msln)* were increased, validating our approach (Figure 5A). As expected the expression of *Nf2* and *Bap1*, tumor suppressors frequently mutated in mesothelioma (Bueno et al., 2016), were significantly decreased during mesothelioma development (Figure 5B). *Nf2* is an upstream regulator of the Hippo signaling cascade (Dong et al., 2007), which prevents nuclear YAP/TAZ localization and activation of transcription enhancers activation domain (TEAD) family members (Zhao et al., 2008). Accordingly, we observed a progressive increase in nuclear YAP localization in parallel to an increase in the proliferation marker Ki-67 in benign growth and tumor tissue from crocidolite-exposed mice (Figure 5C).

**Figure 5.**
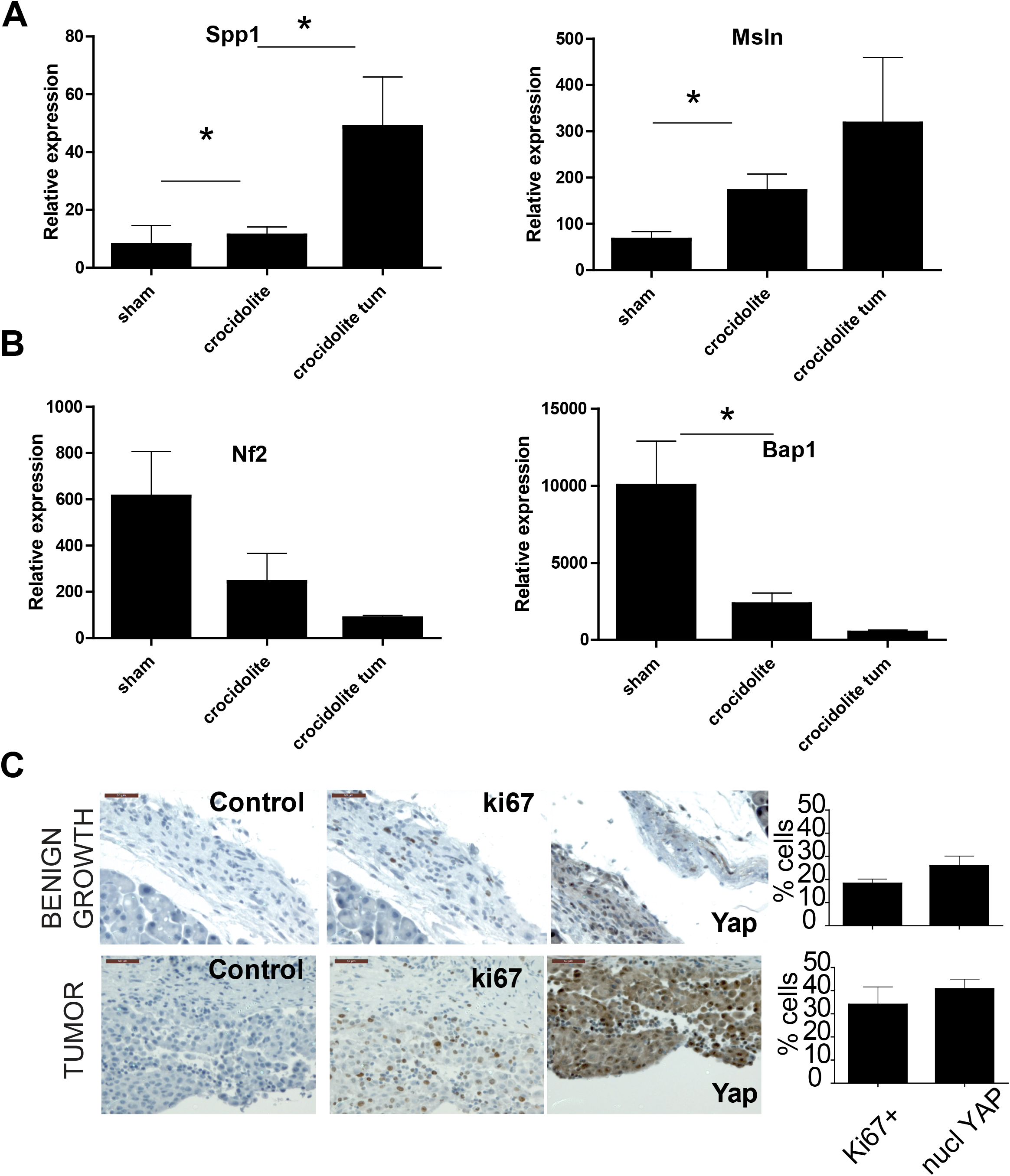
Progressive deregulation of the Hippo pathway during mesothelioma development. **A.** Increased expression of *Spp1* and *Msln* mesothelioma markers in crocidolite-exposed mice validates our approach. qPCR for *Spp1* and *Msln* expression was performed in sham, crocidolite-exposed mice without malign tumor and tumors. Mean±SE. N=5-8 mice, *p< 0.05, Mann-Whitney test. **B.** *Nf2* and *Bap1* tumor suppressors were significantly decreased during mesothelioma development. Gene expression of *Nf2* and *BAP1* were analyzed as in **A.** *p< 0.05, Mann-Whitney test. **C.** Nuclear YAP and ki67 expression in crocidolite-exposed mice. Immunoreactivity for Ki67 and YAP observed in benign growth and in tumors in crocidolite-exposed mice.

Gene set enrichment analysis (Subramanian et al., 2005) of differentially expressed genes indicated that the most significant pathway activated in inflamed tissue compared to sham was activation of interferon-γ (Table S1, p=3.89E-79), while enrichment in epithelial-mesenchymal transition (EMT) in tumors was the most significant pathway in tumors compared to crocidolite-exposed inflamed tissue (Table S2, p=8.92E-81). Many of these genes were downstream of p53 activation, replication stress or YAP activation (Table S3). The number of upregulated genes in the pool of YAP activation pathway signature genes was almost doubled between tumors and inflamed tissue (Figure 6A). One of these genes, *Ctgf*, showed increased expression in tumors, but not in inflamed tissues (Figure 6B), consistent with *Ctgf* expression being synergistically enhanced by TGFβ (Fujii et al., 2012) and enrichment of EMT in tumors.

**Figure 6.**
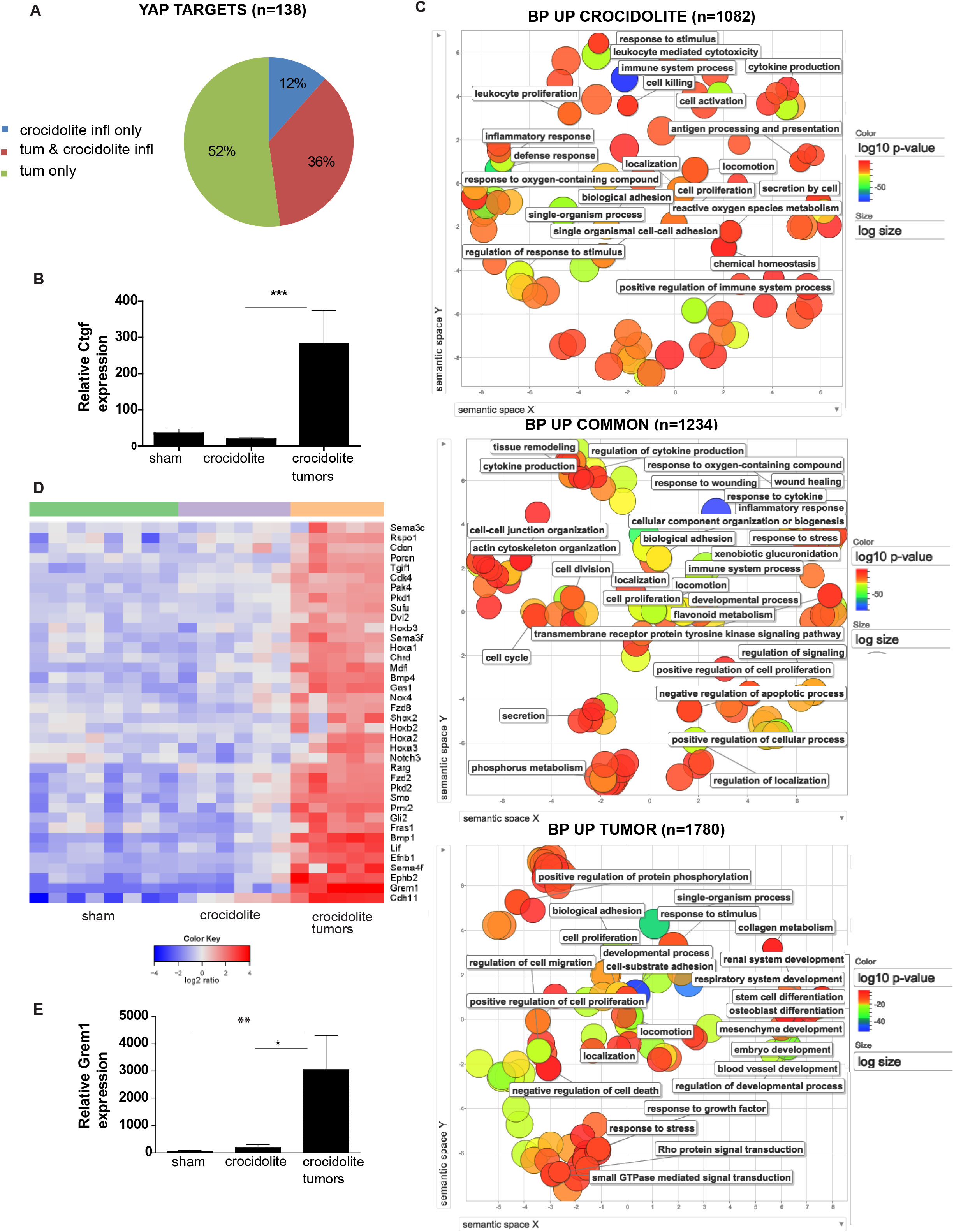
Activation of developmental pathways in mesotheliomagenesis. **A.** Pie chart of significantly increased YAP1 target genes distribution in inflamed tissues vs. tumor. The detailed list of genes is listed in Table 1. **B.** q-pcr of *Ctgf* expression was performed in sham, crocidolite exposed mice without malignant tumor and tumors. Mean±SE. N=5-8 mice. ***p< 0.005, Mann-Whitney test. **C.** The list of genes from each of the three categories of the Venn diagram of Figure 4 were subjected to GO analysis to identify enriched GO terms. The results are shown as REVIGO scatter plots in which similar terms are grouped in a two-dimensional space based on semantic similarity. Each circle indicates a specific GO term and circle sizes are indicative of how many genes are included in each term. Colors indicate the level of significance of enrichment of the GO term. **D.** A curated heatmap was generated for gene sets belonging to categories related to development and that were enriched in tumors. Red represents increased expression. **E.** Overexpression of *Grem1* in tumors was verified by quantitative PCR. Mean±SE. N=5-8 mice. *p< 0.05, **p< 0.01, Mann-Whitney test.

### Activation of developmental genes is predominant in tumors while both inflamed mesothelial tissue and tumor contain Arg1 positive cells

To further explore unique and common genes, we used Gene Ontology Term Finder (http://go.princeton.edu/cgi-bin/GOTermFinder) to find associated GO terms, and *Reduce and Visualize Gene Ontology* (REVIGO) (Supek et al., 2011) to group terms in a two-dimensional space based on semantic similarity (Figure 6C). While biological processes upregulated in crocidolite-exposed tissues included *cytokine production, inflammatory response* and *leukocyte mediated cytotoxicity, cell cycle* and *wound healing* were upregulated in both, crocidolite-exposed non-tumor and tumor tissues. In the latter, Gene Ontology (GO) categories linked several genes to developmental processes. Figure 6D shows a heatmap for selected genes belonging to categories related to development. Ten of the identified 38 genes are part of the Hedgehog pathway and Ephrin/Semaphorin pathways including *Gas1, Cdon, Gli2, Smo, Sufu; Sema4f, Ephb2, Sema3f, Sema3c* and *Efnb1*, consistent with TCGA data showing that these two pathways are part of the top 10 pathways deregulated in human mesothelioma (2014). The Hedgehog signaling pathway is essential during embryonic mesothelial development (Dixit et al., 2013) and is inactive in most adult tissues including mesothelium, but increased Hedgehog signaling is observed in mesothelioma tumors (Shi et al., 2012), especially in the sarcomatoid histotype (Bueno et al., 2016). Five genes (*Fzd8, Fzd2, Dvl2, Porcn, Rspo1*) are part of the *Wnt* pathway and eight genes are part of homeobox families: 5 (*HoxaA1, HoxaA2, HoxA3, HoxB2, HoxB3*) from the Hox family, which defines the morphology of a specific body segment during embryonic patterning and participates in tissue regeneration. The rest of the genes are either part of the BMP/Tgf beta pathway (*Bmp1, Bmp4, Grem1, Chor, Cdh11, Tgfi)*, genes implicated in kidney development (4 genes) or in the *Notch* pathway (2 genes). Few other genes have already been associated with human mesothelioma such as *Lif* (Pass et al., 1995). Analysis of pleural mesothelioma compared to matched normal tissue data (Suraokar et al., 2014) revealed that expression of these genes was also increased in human pleural tumors compared to normal tissue, indicating that activation of these developmental pathways is not species specific. Interestingly, one of the genes with the strongest upregulation φ=2.84E-12, FDR=2.05E-11) in tumors compared with crocidolite-exposed tissue was *Grem1*, a secreted bone morphogenetic protein (BMP) antagonist, which has been recently described to promote epithelial-mesenchymal transition in mesothelioma (Tamminen et al., 2013) (Figure 6E).

Among the genes with the highest upregulation in crocidolite-exposed tissues vs. sham-exposed, and with further upregulation in crocidolite-induced tumor tissue compared to crocidolite-exposed tissue was *Arg1* (p=6.83E-65, FDR=2.98E-61, Figure 7A). Arginase 1 is one of the 2 enzymes, which hydrolyzes L-Arg to urea and L-ornithine, the latter being the main substrate for the production of polyamines that are required for cell cycle progression. Arginase 1 expression was also confirmed in crocidolite-exposed tissue and tumors by immunohistochemistry (Fig 7A). The expression of Arginase 1 is considered to be the hallmark of the “M2”-state macrophage population, which can cause T-cell anergy (Sica and Mantovani, 2012), however it was expressed also in some reactive mesothelial cells which were not stained with the macrophage marker F4/80 (Figure S2). Interestingly the gene signature implicated in the activation of tissue macrophage self-renewal and embryonic stem cells (Soucie et al., 2016), was upregulated in crocidolite-exposed tissues and tumors (Figure 7B), supporting the hypothesis of environment-driven stimulation of pluripotency potential. This observation was reinforced by unbiased oPOSSUM analysis (http://opossum.cisreg.ca/cgi-bin/oPOSSUM3/opossum_mouse_tca) indicating that *Klf4* ranked first among the transcription factors with over-represented binding sites in genes that were upregulated in asbestos-induced inflamed tissue (Z-Score=46.47). Furthermore, curated heatmaps confirmed upregulation of molecules associated with the “M2”-polarized macrophage such as *Csf1, Csfr1* and *Cd163* (Figure S3). Altogether these observations suggest that the increase in the macrophage population resulting from crocidolite exposure is mostly due to “M2”-polarized macrophages.

**Figure 7.**
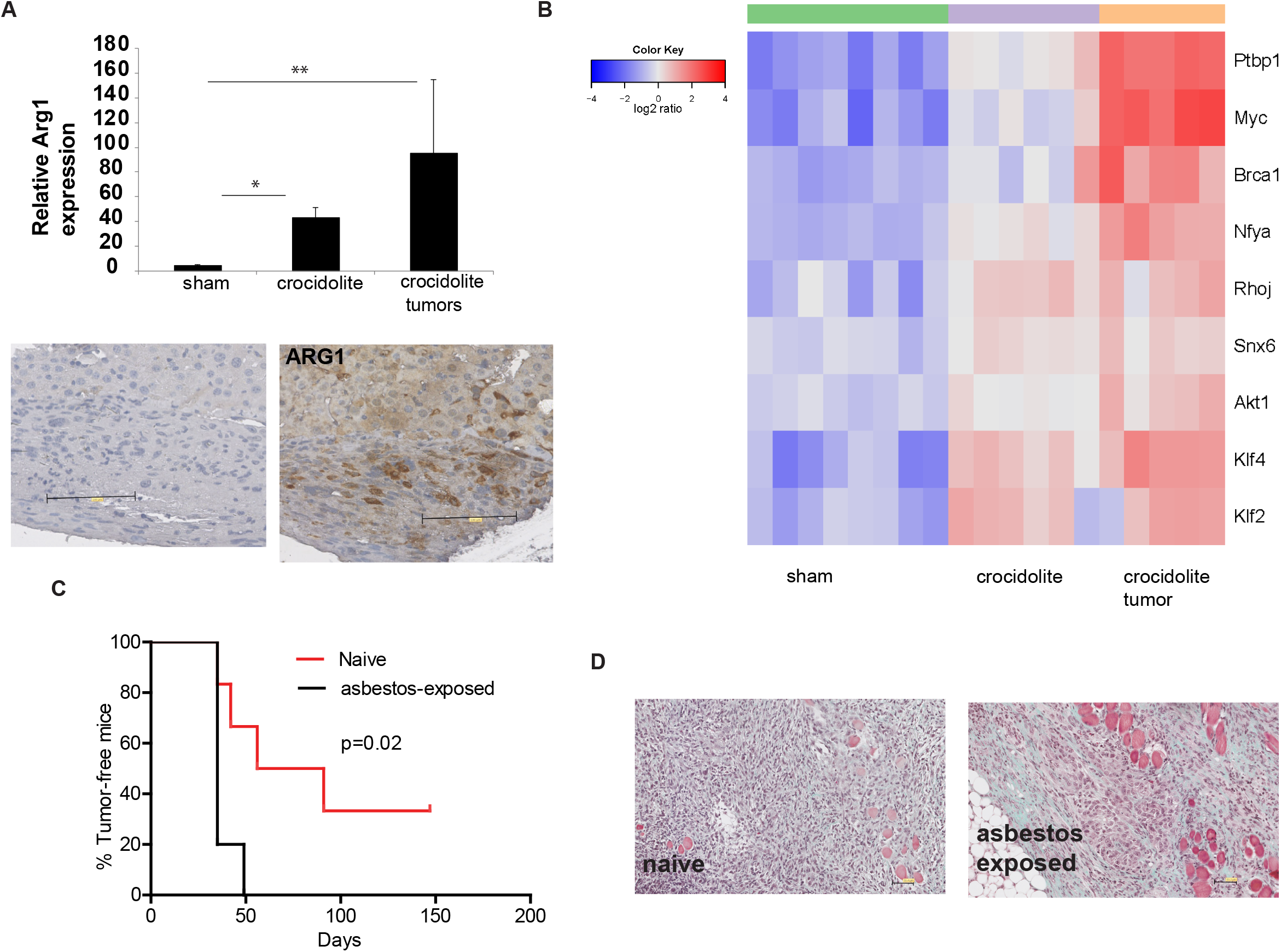
Increase in Arg1 positive cells in both inflamed mesothelial and tumor tissues. **A.** Relative expression of *Arg1* was verified by q-PCR. *Arg1* was highly increased in crocidolite-exposed tissue and expression was maintained in tumors Mean±SE. N=5-8 mice. *p< 0.05, **p< 0.01, Mann-Whitney test compared to sham. Arginase 1 (Arg1)-positive cells were detected in the tumors. **B.** Marker genes for self-renewing macrophages (Soucie et al., 2016) shows increased expression in crocidolite-exposed tissues and tumors. **C.** Kaplan-Meier graph of tumor-free mice survival after challenging with RN5 mesothelioma cells (1 x 10^6^) mice exposed to asbestos that had not developed a tumor until 49 weeks after the first exposure to asbestos vs naïve animals. **D.** Gomorri staining of mesothelioma grown in naïve vs asbestos-exposed mice.

Expression levels of chemokines such as *Ccl2, Ccl5* and *Cxcl-1* were increased in non-tumor tissue from crocidolite-exposed mice (Figure S4), consistent with the observed increase of these signaling molecules in the peritoneal lavage, and even higher levels were measured in tumor tissue. In addition, we observed upregulation of other chemokine receptors such as *Ccr1, Ccr3, Ccr5, Cxcr3, Cxcr4, Cxcr6*, Cx3cr1 and their ligands (Figure S4).

### Inflamed environment favors tumor growth

Finally, in order to investigate whether inflammation observed after asbestos exposure in non-tumor bearing mice would affect tumor growth, tumor-free animals 49 weeks after the first asbestos exposure were challenged with s.c. flank injection of a dose of RN5 syngeneic cells (1 x 10^6^) expected to give rise to tumors only in a fraction of age-matched recipients naïve mice. One mouse exposed to asbestos developed a peritoneal mesothelioma and all the other asbestos-exposed mice developed a s.c. tumor within 50 days while two naïve mice were still tumor-free 150 days after injection of RN5 cells (Figure 7C), supporting the concept of chronic inflammation/mesothelial environment changes predisposing to tumor development. However, no obvious differences in tumor morphology or infiltration were noticed (Figure 7D).

## Discussion

In this study we describe changes in the mesothelial environment and mesothelial tissue during mesothelioma development and provide strong evidence that mesothelioma might be caused by conditions that trigger chronic inflammation (Shalapour and Karin, 2015) where activation of YAP/TAZ signaling, RNA editing and tumor-promoting environment appear prior to tumor development. The RNA editing signature downstream ADAR activity is suggested by A to G mutations at known A to I sites and is consistent with its increased expression and knowledge about *Adar* induction by interferon (George and Samuel, 1999). Decreased expression of *ADARB1* has been observed upon YAP silencing in mesothelioma cells which resulted in decreased cell growth (Mizuno et al., 2012) providing a possible mechanism behind the TCGA data associating high *ADARB1* expression with worst overall survival. RNA editing by ADAR is the most prevalent type of RNA editing and occurs mostly in non-coding regions where inverted repeated sequences are likely to form dsRNA structures, which functions as substrate. It has recently attracted attention as cancer driver for its role in cancer stem cell maintenance, possibly linked also to reduced production of mature miRNA due the impairment of their processing by Drosha after editing of hairpin structures in primitive miRNA (Nishikura, 2016, Jiang et al., 2017). Although we could not evaluate miRNA abundance or mutations due to the method used to extract RNA, it is worth noting that YAP activation has been shown to decrease the activity of Drosha (Mori et al., 2014), therefore RNA editing and YAP/TAZ activation may converge on profound modification of mature miRNA. The progressive activation of YAP/TAZ signaling is possibly associated with extra-cellular matrix (ECM) remodeling since several GO terms induced by crocidolite are associated with this process. In an elegant study investigating how physical/mechanical stimuli are conveyed by ECM stiffness, Dupont et al (Dupont et al., 2011) performed a bioinformatic analysis on genes differentially expressed in mammary epithelial cells grown on ECM of high versus low stiffness. Strikingly, only signatures revealing activation of YAP/TAZ transcriptional regulators emerged as significantly overrepresented in the set of genes regulated by high stiffness. YAP/TAZ signaling is functionally required for differentiation of mesenchymal stem cells induced by ECM stiffness and nuclear YAP/TAZ localization has been associated with response to increased ECM stiffness in several tissues (Musah et al., 2014, Swift et al., 2013). The progressive deregulation of the Hippo pathway associated with YAP/TAZ activation is consistent with previous published results on the role of NF2 in human mesothelioma development (Jongsma et al., 2008), stressing the importance of this signaling pathway in mesothelioma growth. Interestingly, some differences in the profile of YAP/TAZ targets in inflamed tissues *vs*. tumor suggest the existence of modulation of YAP/TAZ activation (Piccolo et al., 2014) including matrix-remodeling in inflamed tissue or epithelial-to-mesenchymal transition cues in tumors. This would be consistent with the observed enrichment of epithelial-to-mesenchymal transition signature and the overexpression of *Grem1* in mice tumors and *Grem1* upregulation in the sarcomatoid histotype of human mesothelioma (Bueno et al., 2016).

Beachy et al. (Beachy et al., 2004) proposed that chronic tissue repair activates stem cell signaling pathways to regenerate the tissue and that oncogenic events may occur due to persistent system stimulation, leading to the formation of a tumor. For the time being, the cell types that proliferate upon exposure to asbestos fibers are not known. Already in the Eighties it has been claimed that mesothelioma originate from the transformation of multipotent connective tissue stem cells that can differentiate into most mesenchymal patterns (Gormley et al., 1980). In MexTag mice where the SV40 T antigen (Tag) is under the control of the mesothelin promoter and which develop mesothelioma tumors upon exposure to asbestos fibers, Tag is not detected in the unexposed mice (Robinson et al., 2010). Mesothelin is a target of YAP activation (Ren et al., 2011) and we observed that mesothelin expression increased in both, inflamed mesothelium and tumors. This is associated with the accumulation of mesothelin-expressing mesothelial precursors in the lavage of asbestos-exposed mice consistent with a previous study, where a mesothelin-expressing population different from macrophages has been described to accumulate in the peritoneal lavage after mesothelium injury (Carmona et al., 2011).

If the mesothelial precursor population has the capacity of self-renewing, it is tempting to speculate that it could accumulate tumor-initiating mutations. In order to determine the contribution of the accumulation of the precursor population, it would be necessary to perform functional studies for example using mesothelin-deficient mice. Mesothelin-deficient mice have a normal phenotype (Bera and Pastan, 2000) and to our knowledge no study has been performed using mesothelin-deficient mice exposed to asbestos to check whether mesothelin deficiency would affect tumor development. However, to address this question it should be kept in mind that the penetrance of the disease after exposure to asbestos fibers in C57BL/6 genetic background is unexpectedly low (10% in this study using Nf2 heterozygotes mice), consistent with other studies (Huaux et al., 2016, Napolitano et al., 2015), therefore further refinements of the model would be necessary before starting such investigation.

In this study we document for the first time the recruitment of a precursor mesothelial population in asbestos-exposed animals. Interestingly, a CD90/Arginase-1 positive mesothelial cell population able to inhibit T cell activation has been previously described within intraperitoneal cells recovered from human samples (Kitayama et al., 2014), therefore it seems possible that mesothelial cells themselves contribute to immunosuppressive signaling.

Although an asbestos signature was present in pre-neoplastic tissues, some mice did not develop a tumor but maintained the predisposition to promote tumor growth highlighting the importance of the environment (niche). The impact of host factors to promote tumor growth is also supported by the observation that less than 50% of human mesothelioma grow in NOD/SCID mice (Tsao et al., 2016), (Felley-Bosco, unpublished), possibly due to an unfavorable growth microenvironment. As mentioned previously, macrophages are recruited to the site of damage after administration of a single dose of asbestos fibers (Moalli et al., 1987) and it has been recently confirmed that this population is the most abundant after intraperitoneal administration of asbestos in C57Bl/6J mice (Huaux et al., 2016). A role for macrophages in asbestos-related chronic inflammation had been suggested in functional studies where “frustrated phagocytosis” induced by exposure of macrophages to asbestos increased the secretion of mature interleukin-1β by activating the complex Nod-like receptor (NLR)-pyrin domain containing 3, Nlrp3, procaspase-1, and the ASC (apoptosis speck-like protein containing a CARD) adapter, which bridges interactions between the former proteins (Dostert et al., 2008). In a later study, although acute interleukin-1β production was significantly decreased in the NLRP3-deficient mice after the administration of asbestos, NLRP3-deficient mice displayed a similar incidence of malignant mesothelioma and survival times as wild-type mice (Chow et al., 2012), ruling out a role for NLRP3 and IL-1β in mesothelioma development, but not for macrophages themselves, which accumulated similarly in wild type and NLRP3-deficient mice.

Besides experimental models, tumor-associated macrophages are a major component of the immune cell infiltration of the tumor microenvironment in mesothelioma patients and the presence of “M2” polarized macrophages is associated with the worst outcome (Cornelissen et al., 2014, Burt et al., 2011). Recent studies have demonstrated that pleural effusions contain “M2” polarized macrophages (Chene et al., 2016, Lievense et al., 2016), which inversely correlated with T cell *in vivo* and suppressed T-cell proliferation *in vitro* (Lievense et al., 2016). This is consistent with recently described recruitment after tissue injury of cavity macrophages, which then become “M2” polarized (Wang and Kubes, 2016). This would also be consistent with the resident macrophage self-renewal signature that we observed in our study.

In conclusion, our comprehensive analysis of loss of homeostasis in the mesothelial environment in asbestos-exposed mice during tumor development suggest a progressive activation of YAP/TAZ-dependent gene transcription, RNA-editing accumulation of single nucleotide variations and a possible role for immunosuppressive macrophages and/or mesothelial precursor cells in tumor development.

## Experimental procedures

### Detailed methods can be found in the Supplemental Experimental Procedures

#### Experimental model

C57Bl/6J and B6;129S2-Nf2^tm1Tyj^/J mice were obtained from Jackson Laboratories (Bar Harbor, Maine). Nf2+/- mice were backcrossed for ≥ 6 generations on a C57Bl/6J genetic background. Unio Internationale Contra Cancrum (UICC)-grade crocidolite asbestos was obtained from SPI Supplies (West Chester, PA). Fibers were suspended in sterile NaCl (0.9%) triturated 10 times through a 22-gauge needle to obtain a homogenous suspension and injected (400 μg/mouse) into 6 to 8 weeks old mice (n=50, 33 male, 17 female), every 3 weeks for a total of eight rounds (i.e. a total of 3.2 mg of crocidolite per mouse). Sham mice (n=37, 19 male, 18 female) were injected with saline. Mice were monitored for tumor formation and sacrificed either 33 weeks after the first crocidolite injection or some animals that had remained tumor-free were used in a challenging experiment 49 weeks after the first exposure to asbestos. RN5 cells (1 x 10^6^), which were derived from one of the tumors (Blum et al., 2015), were injected s.c. and development of tumor growth was recorded.

#### Staining protocol for flow cytometry

The abdominal cavity was washed with 10 ml of phosphate-buffered saline (PBS). Cells obtained by this procedure were pelleted and supernatant was removed and stored at −80°C for later cytokine analysis. Spleens were placed into ice-cold PBS containing 1% FBS. Peripheral blood was drawn from the heart of mice that were immediately euthanized by inhalation of CO2. All samples were passed through 70 μm cell strainer to achieve single cells. ACK lysis buffer (Invitrogen, Carlsbad, CA) was added to spleen samples and allowed to react for at least 15min at room temperature (RT) to lyse red blood cells. After washing thrice with staining buffer, appropriate dilutions (1:50~100) of antibodies or isotype controls were added to each tube, 15min at RT in the dark. Becton Dickinson LSR II Flow Cytometer (San Jose, CA) and FACSDiva™ software were used for data acquisition and FlowJo^TM^ software was used for analysis.

#### Measurement of cytokines/chemokines

Peritoneal lavage fluids were concentrated by ultrafiltration through a low-adsorption polyethersulfonate (PES) membrane (mol. mass. cutoff 3kDa) (concentrator Pierce PES 3K, Thermofisher). The concentration factor was noted for each fluid and used in the calculation of the results. The average concentration factor was 7 with a range from 4 - 10. A Bio-Plex mouse cytokine assay (BioRad) for simultaneous quantification of the concentrations of several signaling molecules (IL-6, IL-10, G-CSF, GM-CSF, IFN-γ, CXCl1, CCL2, CCL5 and VEGF) was run according to the recommended procedure. For each sample, levels of cytokines are reported (in pg/ml) for the non-concentrated peritoneal lavage fluid considering the concentration factor.

#### Library preparation, cluster generation and sequencing

The quantity and quality of the isolated RNA was determined with a Qubit^®^ (1.0) Fluorometer (Life Technologies, California, USA) and a Bioanalyzer 2100 (Agilent, Waldbronn, Germany). The TruSeq Stranded mRNA Sample Prep Kit (Illumina, Inc, California, USA) was used in the succeeding steps. Briefly, total RNA samples (100 ng) were ribosome depleted and then reverse-transcribed into double-stranded cDNA with actinomycin added during first-strand synthesis. The cDNA samples were fragmented, end-repaired and polyadenylated before ligation of TruSeq adapters. The adapters contain the index for multiplexing. Fragments containing TruSeq adapters on both ends were selectively enriched with PCR. The quality and quantity of the enriched libraries were validated using Qubit^®^ (1.0) Fluorometer and the Bioanalyzer 2100 (Agilent, Waldbronn, Germany). The product is a smear with an average fragment size of approximately 360 bp. The libraries were normalized to 10 nM in Tris-HCl 10 mM, pH 8.5 with 0.1% Tween 20.

The TruSeq SR Cluster Kit v3-cBot-HS or TruSeq PE Cluster Kit v3-cBot-HS (Illumina, Inc, California, USA) was used for cluster generation using 8 pM of pooled normalized libraries on the cBOT. Sequencing was performed on the Illumina HiSeq 2000 paired end at 2 X101 bp or single end 100 bp using the TruSeq SBS Kit v3-HS (Illumina, Inc, California, USA). Adapter sequences are available upon request. RNA-seq data are deposited in the European Nucleotide Archive, accession no PRJEB15230.

#### Statistical analysis

Unpaired Student’s t-test, Mann–Whitney U or Kruskal-Wallis test were generally used. Error bars indicate the standard error of the mean. Statistical analysis was performed using Prism 6 (Graphpad)

## Author Contributions

EFB, MdP and BS designed and analyzed all the experiment and wrote the manuscript. LW, LP, WB, EFB and TH contributed to animal exposure to asbestos and samples collection. LW, TH, WB, EFB and LP contributed to experiments using syngeneic mesothelioma cells. LW performed flow cytometry experiments. EFB performed the extraction of RNA, CA performed RNA-seq and HR and EFB analyzed the gene expression data. VSB quantified signaling molecules in peritoneal lavage. BV assessed histopathology.

## Acknowledgments

The project is financed by the Swiss National Science Foundation Sinergia grant CRSII3 #147697. We are grateful to Dr. Marie-Claude Jaurand and Dr. Maries van den Broek for critical reading of the manuscript. We are also grateful to Manuel Ronner, Gabriela Ziltener, Janine Worthmüller, Simone Eichenberger, Marlène Sanchez for skillful assistance. We also thank the Princess Margaret Hospital Foundation and the Toronto General and Western Hospital Foundation to MdP, the Stiftung für Angewandte Krebsforschung to EFB. We thank Dr. Roman Briskine for running the SNV analysis workflow.

The authors declare no conflicts of interest.

